# Differential requirement for BRCA1-BARD1 E3 ubiquitin ligase activity in DNA damage repair and meiosis in the *Caenorhabditis elegans* germ line

**DOI:** 10.1101/2022.10.02.510574

**Authors:** Qianyan Li, Arshdeep Kaur, Kyoko Okada, Richard J. McKenney, JoAnne Engebrecht

**Affiliations:** Department of Molecular and Cellular Biology, University of California Davis, Davis, CA 95616, USA; Biochemistry, Molecular, Cellular and Developmental Biology Graduate Group, University of California Davis, Davis, CA 95616, USA

## Abstract

The tumor suppressor BRCA1-BARD1 complex functions in many cellular processes; of critical importance to its tumor suppressor function is its role in genome integrity. Although RING E3 ubiquitin ligase activity is the only known enzymatic activity of the complex, the *in vivo* requirement for BRCA1-BARD1 E3 ubiquitin ligase activity has been controversial. Here we probe the role of BRCA1-BARD1 E3 ubiquitin ligase activity *in vivo* using *C. elegans*. Genetic, cell biological, and biochemical analyses of mutants defective for E3 ligase activity reveal both E3 ligase-dependent and independent functions of the complex in the context of DNA damage repair and meiosis. We show that E3 ligase activity is essential for BRCA1-BARD1 to concentrate at both DNA damage and recombination sites in meiotic germ cells, but not at DNA damage sites in proliferating germ cells. While BRCA1 alone is capable of monoubiquitylation, BARD1 is required with BRCA1 to promote polyubiquitylation. We find that the requirement for E3 ligase activity and BARD1 in DNA damage signaling and repair can be partially alleviated by driving the nuclear accumulation and self-association of BRCA1. Our data suggest that in addition to E3 ligase activity, BRC-1 serves a structural role for DNA damage signaling and repair while BRD-1 plays an accessory role to enhance BRC-1 function.

**Author Summary:** BRCA1-BARD1 is a E3 ubiquitin ligase, which modifies proteins by the addition of the small protein ubiquitin. While mutations that disrupt E3 ligase activity and stability of the BRCA1-BARD1 complex lead to a predisposition for breast and ovarian cancer, the specific requirement for E3 ligase activity in tumor suppression is not known. Here we probe the function of E3 ligase activity and BARD1 in the maintenance of genome integrity by engineering point mutations that disrupt E3 ligase activity in *C. elegans* BRCA1 as well as a null mutation in BARD1. We find that while E3 ligase activity is important for genome integrity, the complex plays additional roles besides ubiquitylating proteins. Further, our data suggest that BRCA1 is the key functional unit of the complex while BARD1 is an accessory partner that enhances BRCA1’s function. These findings may help explain why there is a higher prevalence of cancer-causing mutations in BRCA1 compared to BARD1.

## Introduction

<u>BR</u>east <u>CA</u>ncer susceptibility gene 1 (BRCA1) and its obligate partner BARD1 (<u>B</u>RCA1 <u>A</u>ssociated <u>R</u>ING <u>D</u>omain protein 1) are RING domain-containing proteins, which when mutated are linked to elevated incidence of breast and ovarian cancer (1-6). The BRCA1-BARD1 complex functions in a myriad of cellular processes, including DNA damage repair, replication, checkpoint signaling, meiosis, chromatin dynamics, centrosome amplification, metabolism, and transcriptional and translational regulation (7-13). BRCA1-BARD1 regulates these pathways presumably through ubiquitylation of substrates via its RING domains, which function as an E3 ubiquitin ligase. BRCA1 specifically interacts with E2-conjugating enzymes for ubiquitin transfer, while BARD1 greatly stimulates the E3 ligase activity of BRCA1 (14, 15).

Multiple potential BRCA1-BARD1 substrates have been identified; however, the physiological significance of most of these substrates is currently unknown (16). One well established substrate in the context of DNA damage signaling and transcriptional regulation is histone H2A (17-19). Recent structural and molecular studies have led to mechanistic insight into recruitment of the complex to DNA damage sites and subsequent ubiquitylation of histone H2A. These studies have highlighted the targeting role of BARD1 to nucleosomes, where ubiquitylation of H2A by the complex promotes repair of DNA double strand breaks (DSBs). This most likely occurs by blocking recruitment of 53BP1, which promotes error-prone non-homologous end joining at the expense of homologous recombination (20-22). However, the full spectrum of substrates and their relationship to regulation of different cellular processes are currently not known.

The role of BRCA1-BARD1 in DNA damage repair has been linked to its tumor suppressor function. Early studies suggested that BRCA1-BARD1 E3 ligase activity was not essential for either recombinational repair or tumor suppression. This conclusion was based on the analysis of a single isoleucine to alanine mutation at amino acid 26 (I26A) in the BRCA1 RING domain that abrogates its E3 ligase activity *in vitro* but maintains the stability of the BRCA1-BARD1 heterodimer, unlike many cancer-causing mutations that impair both E3 ligase activity and heterodimer stability (14, 23-26). Mice expressing the BRCA1^I26A^ mutant protein were not prone to tumor formation and mutant cells were proficient for homology-directed repair of DSBs, suggesting that E3 ligase activity is not essential for tumor suppressor function (27, 28). In depth biochemical analyses, however, have shown that the BRCA1^I26A^ mutant still exhibits residual E3 ligase activity when paired with a subset of E2 ubiquitin-conjugating enzymes in *in vitro* ubiquitin transfer assays. Mutation of two additional residues (leucine 63 and lysine 65 changed to alanines) within the BRCA1 RING domain in combination with I26A are required to completely abrogate E3 ligase activity *in vitro* without compromising the structural integrity of the complex (29). These results suggest that BRCA1 harboring all three mutations is a true ligase dead mutant; however, the phenotypic consequence of this triple mutation has not been analyzed.

To define the requirement for E3 ligase activity *in vivo*, we focused on the *C. elegans* BRCA1 and BARD1 orthologs, BRC-1 and BRD-1. Previous analyses revealed that *C. elegans* BRC-1-BRD-1 is a functional E3 ubiquitin ligase that plays roles in DNA damage repair and meiotic recombination (30-40), as well as regulation of heterochromatin during embryogenesis (41). Additionally, a recent study reported a role for BRC-1-BRD-1 in post-mitotic axon regeneration (42), consistent with the complex playing multiple roles *in vivo*. Here we analyzed the requirement for BRC-1-BRD-1 E3 ubiquitin ligase activity by generating worms expressing BRC-1 mutant proteins containing the corresponding single (I23A) and triple (I23A, I59A, R61A) mutations based on modeling with human BRCA1. We found both E3 ligase-dependent and independent functions of BRC-1-BRD-1 in the context of DNA damage repair and meiosis. Intriguingly, E3 ligase activity and BRD-1 function can be partially bypassed by independently driving nuclear accumulation and self-association of BRC-1. Our data suggest that in addition to E3 ligase activity, BRC-1 serves a structural role for DNA damage signaling and repair while BRD-1 plays an accessory role to enhance BRC-1 function.

## Results

### *brc-1(triA)* exhibits a more severe phenotype than *brc-1(I23A)*

The *C. elegans* orthologs of BRCA1 and BARD1 are structurally conserved with the same key domains as the human proteins: both BRC-1 and BRD-1 contain an N-terminal RING domain and C-terminal BRCT repeat domains (31). The RING domains specify E3 ubiquitin ligase activity, while BRCT domains are phospho-protein interaction modules. Sequence alignment between human BRCA1 and *C. elegans* BRC-1 RING domains reveals that residues essential for E3 ligase activity in human BRCA1 (isoleucine 26, leucine 63, and lysine 65) correspond to amino acids isoleucine 23, isoleucine 59 and arginine 61 in *C. elegans* BRC-1 (Fig 1A). While not identical, these amino acids have similar chemical properties in terms of hydrophobicity and charge. To confirm that these BRC-1 residues structurally align with the human residues critical for ubiquitin transfer, we used AlphaFold to predict the structure of *C. elegans* BRC-1 RING domain, which was superimposed onto the NMR structure of the human BRCA1 RING domain (Fig 1B) (14, 43, 44). The predicted structure overlay is consistent with the sequence alignment in that isoleucine 23, isoleucine 59 and arginine 61 in BRC-1 are the structural counterparts of isoleucine 26, leucine 63 and lysine 65 in human BRCA1.

**Fig 1.**
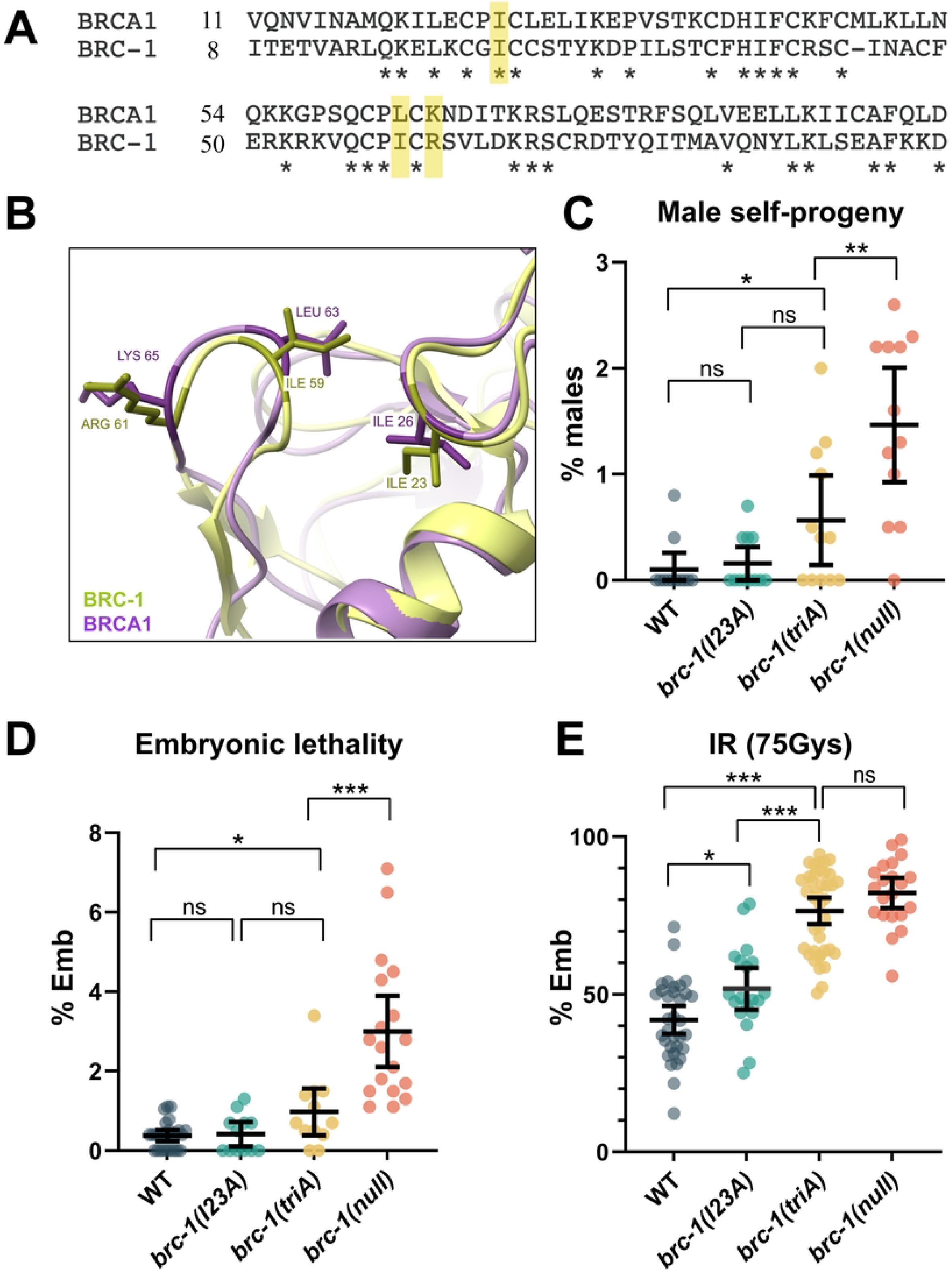
Mutation of three key amino acids in the BRC-1 RING domain leads to a more severe phenotype than the single I23A mutation. (A) Sequence alignment reveals that amino acids isoleucine 23, isoleucine 59 and arginine 61 in *C. elegans* BRC-1 RING domain correspond to isoleucine 26, leucine 63 and lysine 65 in human BRCA1 RING domain (yellow). (B) Structure of BRC-1 RING domain (green) predicted by AlphaFold superimposed onto the NMR structure of human BRCA1 RING domain (purple) showing the three amino acids occupy the same physical position. (C) Male self-progeny, (D) embryonic lethality (Emb), and (E) embryonic lethality in the presence of 75Gys IR were examined in wild type, *brc-1(I23A), brc-1(triA)*, and *brc-1(null)* animals. Number of animals examined in (C): n=12 for all genotypes; (D): WT n=26; *brc-1(I23A)* n=12; *brc-1(triA)* n=12; *brc-1(null)* n=18; (E) WT n=38; *brc-1(I23A)* n=19; *brc-1(triA)* n=39; *brc-1(null)* n=21. *** p < 0.001; ** p < 0.01; * p < 0.05; ns = not significant by Mann-Whitney.

To probe the *in vivo* function of BRC-1 E3 ligase activity, we generated *C. elegans* mutants *brc-1(I23A)* [isoleucine 23 mutated to alanine] and *brc-1(triA)* [isoleucine 23, isoleucine 59, arginine 61 mutated to alanines] at the endogenous *brc-1* locus using CRISPR-Cas9 genome editing and analyzed the mutant phenotypes with respect to meiosis and DNA damage repair. *brc-1* and *brd-1* mutants produce slightly elevated levels of male self-progeny (X0), a readout of meiotic X chromosome nondisjunction, have low levels of embryonic lethality under standard growth conditions, but display high levels of embryonic lethality after exposure to γ-irradiation (IR), which induces DNA DSBs (31, 32, 34). For both male self-progeny and embryonic lethality under standard growth conditions, the *brc-1(I23A)* mutant produced similar levels to wild type, while *brc-1(triA)* worms gave rise to elevated levels compared to wild type but not to the extent of the *brc-1(null)* mutant (34) (Fig 1C, D). Following exposure to 75Gys of IR, *brc-1(I23A)* displayed higher levels of embryonic lethality compared to wild type, while *brc-1(triA)* produced inviable progeny at levels comparable to those observed in the *brc-1(null)* mutant, suggesting that E3 ligase activity is important when DNA damage is present.

While BRC-1-BRD-1 plays only a subtle role in an otherwise wild-type meiosis as evidenced by the low levels of male self-progeny and embryonic lethality (Fig 1C, D), we previously showed that the BRC-1-BRD-1 complex plays a critical role when chromosome synapsis and crossover formation are perturbed by mutation of either pairing center proteins, which are required for pairing and synapsis of homologous chromosomes, or components of the synaptonemal complex (SC), the meiosis-specific protein structure that stabilizes homologous chromosome associations (33, 34). To examine the phenotypic consequence of impairing BRC-1-BRD-1 E3 ligase activity when meiosis is perturbed, we monitored embryonic lethality in the different *brc-1* mutants in combination with mutation of ZIM-1, a zinc finger pairing center protein that mediates the pairing and synapsis of chromosomes *II* and *III* (45). *zim-1* mutants produce 60-70% inviable progeny due to random segregation of chromosomes *II* and *III* in meiosis resulting in the formation of aneuploid gametes. We observed a progressive increase in embryonic lethality in *brc-1(I23A); zim-1, brc-1(triA); zim-1*, and *brc-1(null); zim-1* mutants, consistent with our previous observation that *brc-1(triA)* is more severely impaired for function than *brc-1(I23A)*. These results also suggest that neither *brc-1(I23A)* nor *brc-1(triA)* are null alleles (Fig 2A).

**Fig 2.**
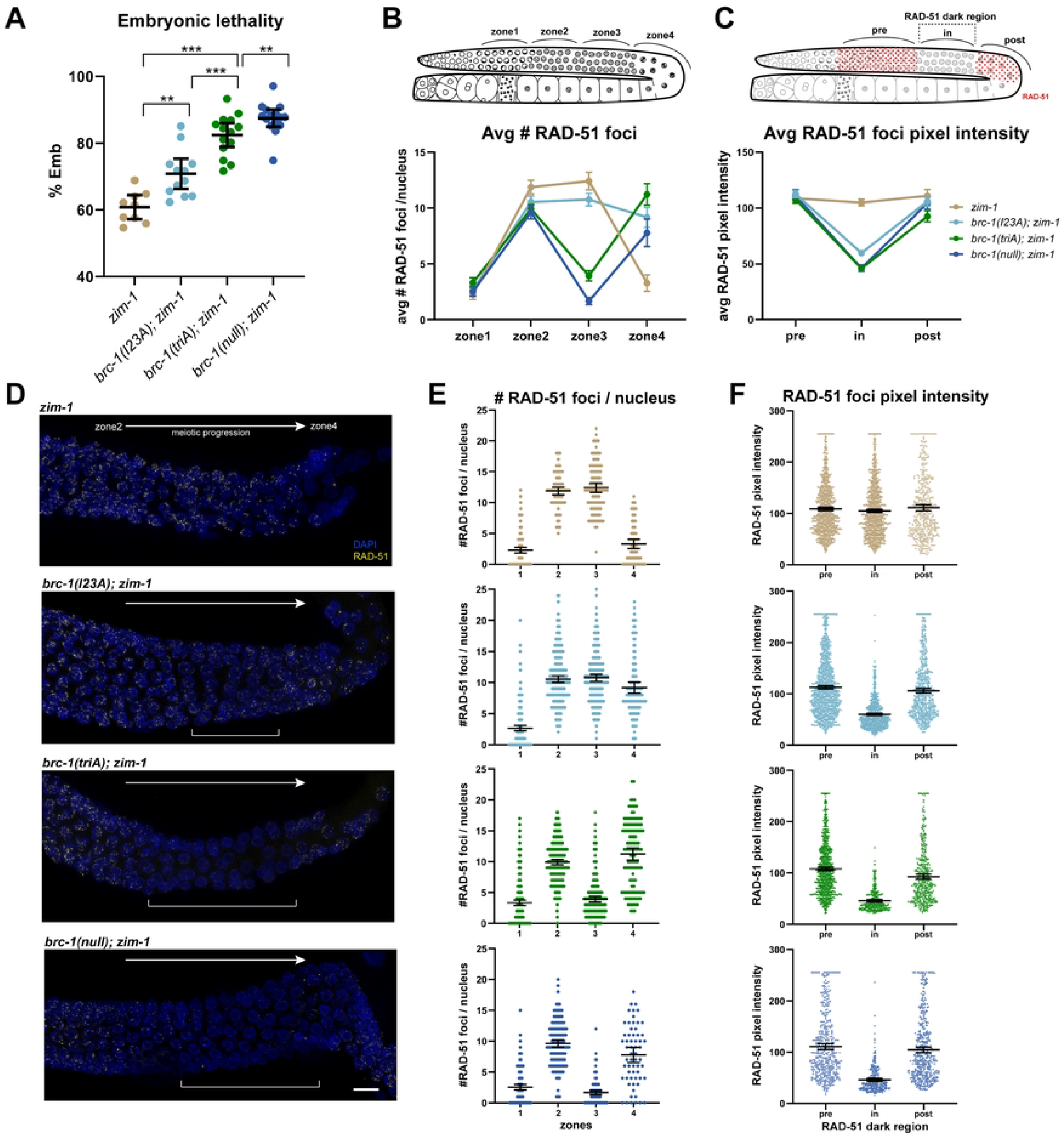
*brc-1(I23A)* and *brc-1(triA)* mutants show differential defects in promoting progeny viability and RAD-51 filament stabilization in the *zim-1* mutant. **(A)** Embryonic lethality of *brc-1* mutants in the *zim-1* mutant background. Embryonic lethality of *brc-1(triA); zim-1* mutant (n=14) is intermediate between *brc-1(I23A); zim-1* (n=12) and *brc-1(null); zim-1* mutants (n=15); *zim-1* (n=9)*** p < 0.001; ** p < 0.01 Mann-Whitney. (B) Cartoon of gonad indicating the zones analyzed for RAD-51 foci numbers across the meiotic region. Graph depicts the average number of RAD-51 foci per nucleus quantified per zone from three germ lines of indicated genotypes. RAD-51 foci number only modestly declines in zone 3 in the *brc-1(I23A); zim-1* mutant. (C) Cartoon of gonad indicating regions analyzed for RAD-51 foci pixel intensity. Graph shows average pixel intensity of RAD-51 foci from pre, in and post RAD-51 dark region in three germ lines of the indicated genotypes. *brc-1(I23A); zim-1* contains nuclei with reduced RAD-51 foci intensity in the dark region but not to the extent of *brc-1(triA); zim-1* and *brc-1(null); zim-1* mutants. (D) Images showing part of the germ line from early/mid-pachytene (zone 2) to diplotene (zone 4) immunolabeled with RAD-51 antibody (yellow) and counterstained with DAPI (blue). Brackets indicate the presence and location of RAD-51 dark region in the mutant germ lines, which is not as pronounced in the *brc-1(I23A); zim-1* mutant. Scale bar = 10μm (E) Scatter plot of number of RAD-51 foci per nucleus across the four zones. (F) Scatter plot of RAD-51 foci pixel intensity from pre, in and post RAD-51 dark regions in the germ lines. Mean and 95% CI are indicated for all data sets; statistical comparisons between genotypes are shown in Supplemental Table 3.

In addition to enhancing embryonic viability, BRC-1 and BRD-1 stabilize the RAD-51 filament at mid to late pachytene in the *zim-1* mutant (33, 34). The RAD-51 recombinase assembles on resected single strand DNA at DSBs and is essential for homology search and strand invasion during homologous recombination (46-48). In mutants where crossover formation is blocked by defects in pairing or synapsis (e.g., *zim-1*), RAD-51 filaments, visualized as nuclear foci by immunostaining, are extended into late pachytene (34, 48, 49). Removal of BRC-1 in this context results in a “RAD-51 dark region” at mid to late pachytene due to a defect in RAD-51 filament stability (34) (Fig 2C, D). This is manifested in a reduction in both RAD-51 foci numbers (zone 3) and signal intensity (in dark region), followed by an increase in RAD-51 foci numbers (zone 4) in the *brc-1(null); zim-1* mutant (Fig 2B-F). To determine whether BRC-1 E3 ligase activity is required for RAD-51 stabilization when meiosis is perturbed, we monitored RAD-51 foci number and signal intensity in the different *brc-1; zim-1* mutants (Fig 2B). As with the increasing severity of embryonic lethality in the putative E3 ligase dead alleles in *zim-1* mutants, impairment of RAD-51 filament stability also showed increasing severity in the mutants (Fig 2D). In zones 2 and 3, *brc-1(I23A); zim-1* had slightly reduced numbers of RAD-51 foci compared to *zim-1*, but significantly higher numbers than observed in *brc-1(null); zim-1*. More foci were observed in zone 4 in *brc-1(I23A); zim-1* than *zim-1* alone (Fig 2B, E; Supplemental Table 3). Additionally, average RAD-51 foci intensity in the *brc-1(I23A); zim-1* was significantly reduced in the RAD-51 dark region compared to *zim-1*, but not as reduced as in the *brc-1(null); zim-1* mutant (“in”; Fig 2C, F; Supplemental Table 3). These findings suggest that *brc-1(I23A)* has a partial defect in RAD-51 filament stabilization. In contrast to *brc-1(I23A); zim-1, brc-1(triA); zim-1* showed a severe reduction in the average number of RAD-51 foci in zone 3, although not to the same extent as in *brc-1(null); zim-1*. There was also a significant reduction in average RAD-51 foci intensity in the RAD-51 dark region in *brc-1(triA); zim-1* comparable to that observed in the *brc-1(null); zim-1* mutant (“in”; Fig 2B-F; Supplemental Table 3). Taken together, *brc-1(I23A)* has a weak phenotype, and *brc-1(triA)* is more severe, but still less severe compared to the *brc-1(null)*, suggesting that while E3 ligase activity of BRC-1 is important in the context of defective meiosis, the presence and structural integrity of the complex is critical.

### BRC-1^I23A^ and BRC-1^triA^ are impaired for E3 ubiquitin ligase activity *in vitro*

To verify that BRC-1^I23A^ and BRC-1^triA^ are impaired for E3 ubiquitin ligase activity, we expressed and purified a chimeric form of the RING domains of BRC-1 and BRD-1 (BRD-1-BRC-1) in *E. coli*, modeled after studies of the human complex (23) (Fig 3A; Supplemental Fig 1A). The BRD-1-BRC-1 chimera was incubated in the presence of human UBE1 E1 activating enzyme (50% identical to *C. elegans* E1 UBA-1), UbcH5c E2 conjugating enzyme (94% identical to *C. elegans* E2 UBC-2), HA-ubiquitin (99% identical to *C. elegans* ubiquitin), and ATP and auto-ubiquitylation of the chimera was used as a readout for E3 ubiquitin ligase activity. Visualization with anti-HA antibodies revealed a characteristic ladder of bands due to the incorporation of multiple HA-ubiquitins into the chimera in the presence of ATP, indicating robust auto-polyubiquitylation catalyzed by the BRD-1-BRC-1 chimera (Fig 3B).

**Fig 3.**
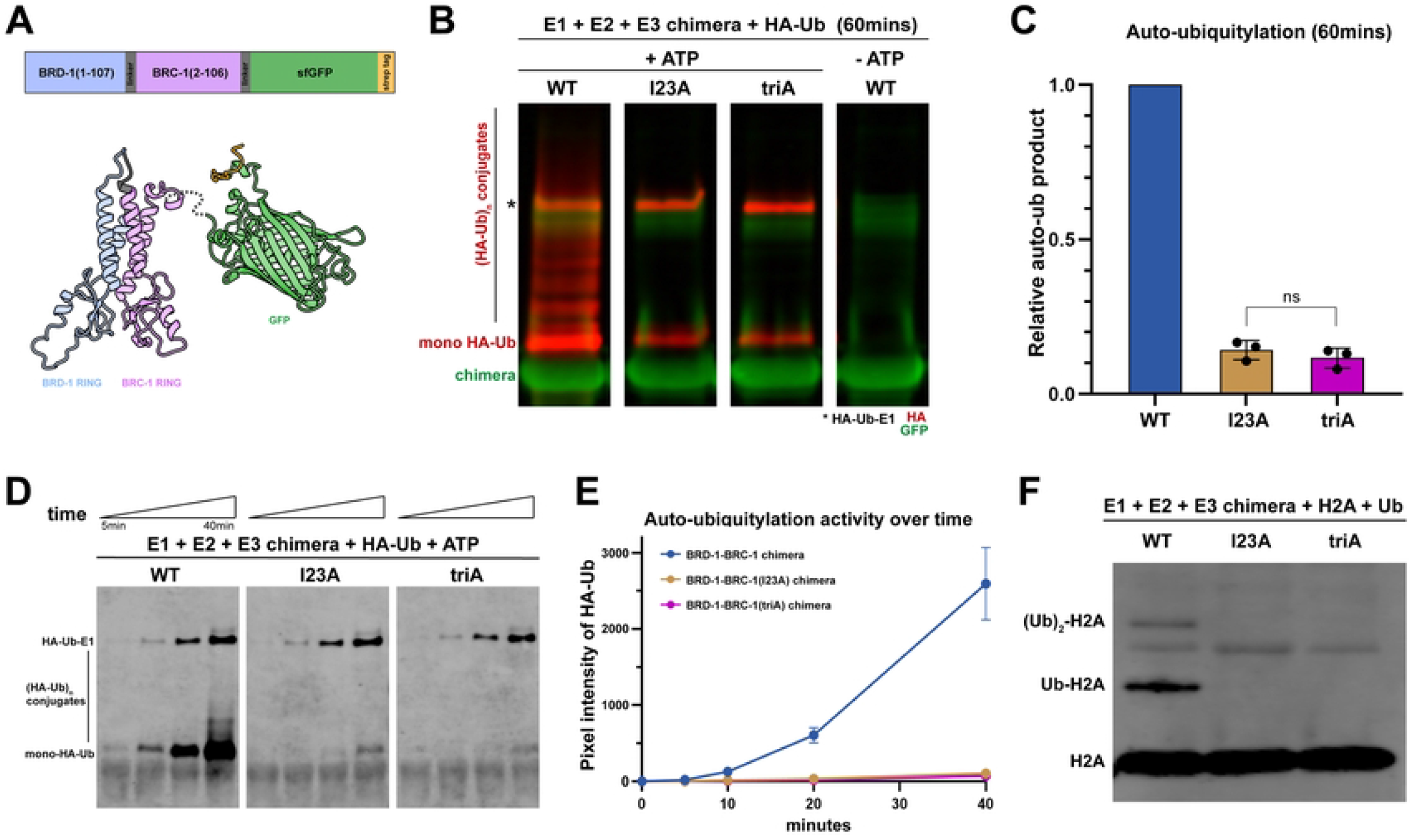
BRD-1-BRC-1^I23A^ and BRD-1-BRC-1^triA^ chimeras are defective for E3 ubiquitin ligase activity *in vitro*. (A) Construct and model based on AlphaFold of BRD-1-BRC-1 chimera: N-terminal BRD-1 RING domain (amino acids 1-107; blue), GGSGG linker (grey) and the BRC-1 RING domain (amino acids 2-106; purple) are connected to a superfold GFP (green) and strep II tag (orange) at the C terminus. Mutant chimera proteins contain either the single I23A or the I23A I59A R61A triple mutations (triA) in the BRC-1 RING. (B) Immunoblot showing auto-ubiquitylation (anti-HA-Ub, red) of BRD-1-BRC-1 chimera (anti-GFP, green) when incubated with E1, E2, HA-Ub and ATP for 60mins. *E1 incorporates HA-Ub (HA-Ub-E1) independently of E2 or E3s. Wild-type chimera promotes the formation of both auto-mono (mono HA-Ub) and polyubiquitylated (HA-Ub_n_) conjugates while only reduced levels of auto-monoubiquitylated BRD-1-BRC-1 were present in mutant chimera reactions. (C) Quantification of total HA-Ub signal at the end of 60mins showed that I23A and triA chimeras produced an average of 14% and 12% of total ubiquitylation, respectively, as compared to the wild-type chimera. The difference between I23A and triA is not significant (ns) by Student T test, p = 0.55. (D) Time-course experiment to compare the kinetics of E3 ligase activity of the wild-type and mutant chimeras. Immunoblot showing HA-Ub signal at 5, 10, 20, and 40mins after the respective chimera was incubated with E1, E2, HA-Ub and ATP. (E) Quantification of HA-Ub signals plotted against time in wild-type and mutant chimeras (At 40mins: I23A = 0.041±0.013, triA = 0.028±0.016 of wild-type auto-ubiquitylation). (F) Immunoblot of *C. elegans* ubiquitin incorporation into human histone H2A (anti-H2A) by WT and mutant chimeras; only WT was able to transfer ubiquitin to histone H2A protein to generate mono (Ub-H2A) and di ((Ub)_2_-H2A) ubiquitylation, but no ubiquitin incorporation into H2A was observed with either the I23A or triA mutant chimeras.

We next expressed and purified mutant chimeras harboring the I23A or triA (I23A, I59A, R61A) mutations (S1A Fig) and performed the auto-ubiquitylation assay. We observed a significant reduction in the incorporation of HA-ubiquitin into the mutant complexes. While no polyubiquitylation was observed, there was reduced but detectable monoubiquitylation of both I23A and triA chimeras by an end point assay (I23A = 14%, triA = 12% of wild-type auto-ubiquitylation; Fig 3C). Time course analyses with decreased concentrations of reaction components revealed a significant reduction in ubiquitin transfer by the I23A and triA chimeras. After 40 mins the I23A and triA chimeras showed only 4.1% and 2.8% of total ubiquitin incorporation as compared to the wild-type chimera (Fig 3D, E). While it did not reach statistical significance, the triA chimera showed consistently lower auto-monoubiquitylation than the I23A chimera. The physiological relevance of the residual auto-monoubiquitylation observed in the mutant chimeras is not clear as it was also observed in reactions lacking the E2 conjugating enzyme (S1B Fig).

Histone H2A is a known physiological substrate of human BRCA1-BARD1 (17, 19). We next determined whether the wild-type and mutant chimeras could catalyze the incorporation of HA-ubiquitin into human histone H2A (90% identical to *C. elegans* H2A). Using antibodies directed against histone H2A or HA, we observed incorporation of mono and di-ubiquitin into H2A in the wild-type reaction; however, no ubiquitin incorporation into H2A was detected with either of the mutant chimeras (Fig 3F; S1C Fig). From these experiments we conclude that BRC-1 harboring the I23A and the triA mutations are both significantly impaired for E3 ubiquitin ligase activity *in vitro*.

### Nuclear accumulation and BRC-1-BRD-1 interaction are differentially affected by BRC-1^I23A^ and BRC-1^triA^

The finding that the *brc-1(I23A)* mutant had considerably weaker phenotypes in DNA damage repair and meiosis compared to the *brc-1(triA)* mutant but displayed similar impairment in E3 ubiquitin ligase activity *in vitro*, led us to examine the consequence of these mutations in more detail. We first monitored the localization of the mutant complexes using antibodies directed against BRD-1(35). BRC-1 and BRD-1 are mutually dependent for localization and are enriched in germ cell nuclei; in mitotic and early meiotic germ cells the complex is observed diffusely on chromatin and in foci. As meiosis progresses BRC-1-BRD-1 becomes associated with the SC and is then restricted to six small stretches on the six pairs of homologous chromosomes defined by the single crossover site (32, 34, 35) (Fig 4A). In the *brc-1(I23A)* mutant BRD-1 displayed a similar localization pattern as wild type, although the intensity of the signal was weaker and not as concentrated in the nucleus (nuclear enrichment in *brc-1(I23A)* was 79% of wild type; Fig 4A, B). Nuclear accumulation of BRD-1 was further impaired in the *brc-1(triA)* mutant in proliferating germ cells through mid pachytene, where the protein was enriched in the cytoplasm relative to the nucleus (Fig 4A, B). At late pachytene and diakinesis in the *brc-1(triA)* mutant, BRD-1 was observed in short stretches in the nucleus, in addition to the cytoplasmic signal (Fig 4A). A similar localization pattern was observed by live cell imaging in the corresponding *brc-1* mutants expressing BRD-1::GFP at the endogenous locus (34) (S2A Fig). These results suggest that nuclear accumulation of the complex is impaired in the E3 ligase defective mutants. Significantly less BRD-1 accumulates in the nucleus in *brc-1(triA)* compared to *brc-1(I23A)*, and this difference likely contributes to the increased severity of the *brc-1(triA)* mutant observed *in vivo*.

**Fig 4.**
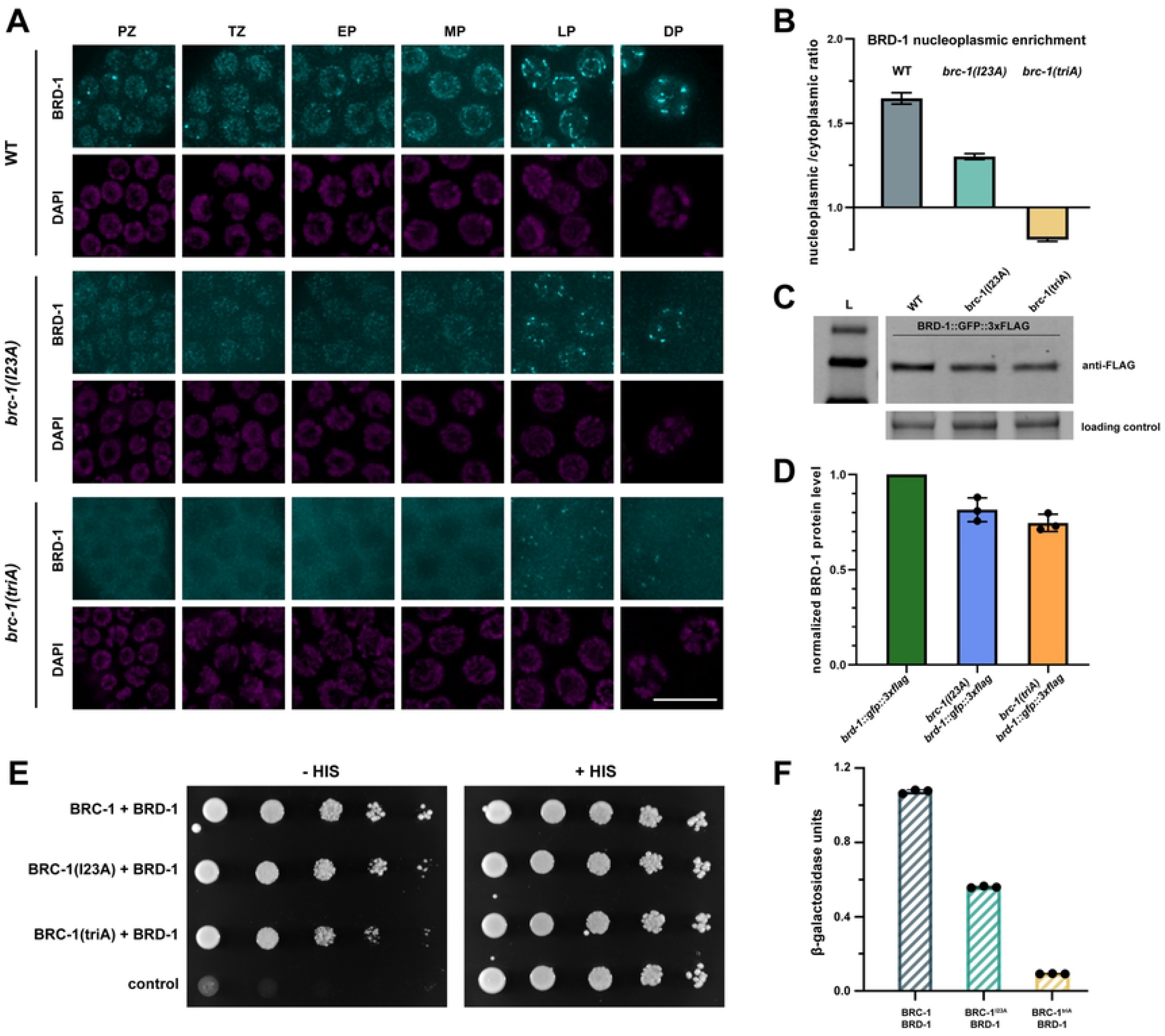
Nuclear accumulation and BRC-1-BRD-1 interaction are differentially affected by BRC-1^I23A^ and BRC-1^triA^ mutations. (A) Images of germline nuclei showing BRD-1 immunolabeling (cyan) by anti-BRD-1 antibodies and DAPI staining to visualize DNA (magenta). PZ = proliferative zone; TZ = transition zone; EP = early pachytene; MP = mid pachytene; LP = late pachytene; DP = diplotene stages in the germ line. Scale bar = 10μm. (B) Graph shows nucleoplasmic to cytoplasmic ratio of BRD-1 signal. (C) Immunoblot of BRD-1::GFP::3xFLAG from whole worm extracts. BRD-1::GFP::3xFLAG migrates slower than its predicted size (112 kDa); molecular weight standards = 170, 130, 100 kDa, respectively. (D) Quantification of BRD-1::GFP::3xFLAG steady state levels in the *brc-1* mutants normalized to the wild type from 3 independent experiments. (E) Yeast two-hybrid interaction between BRC-1 and BRD-1 as measured by growth on medium lacking histidine (-HIS) with +HIS as control. (F) Relative β-galactosidase activity assay showing reduced interaction between mutant BRC-1 (I23A and triA) and BRD-1 in corresponding yeast strains.

Given the reduced signal of BRD-1 observed by immunofluorescence in the *brc-1(I23A)* and *brc-1(triA)* mutants, we next examined steady state protein levels by immunoblot analysis. For these experiments we used worms expressing BRD-1::GFP, which also contain 3 copies of the FLAG epitope, and monitored protein levels using anti-FLAG antibodies. We observed a modest reduction of BRD-1 steady state levels in the *brc-1(I23A)* and *brc-1(triA)* mutants compared to wild type (*brc-1(I23A) brd-1::gfp* = 81.5 ± 6.2% and *brc-1(triA) brd-1::gfp* = 74.6 ± 4.5% of *brd-1::gfp*) (Fig 4C, D). However, there was no significant difference between the two mutants (p=0.19), suggesting that the difference in phenotypes observed *in vivo* is not a consequence of altered steady state protein levels in the mutants, but likely reflects a change in subcellular distribution.

It has been reported that mutations of either I26 or I26, L63, K65 do not alter the interaction between human BRCA1 and BARD1 (29). To determine whether this is also the case for the *C. elegans* orthologs, we examined interaction between full length BRC-1 and BRD-1 using the yeast two-hybrid system, which has previously been used to demonstrate interaction between these two proteins (31). As expected, an interaction was detected between wild-type BRC-1 and BRD-1 using *his3* expression as a reporter by monitoring growth on medium lacking histidine. We observed slightly less growth on medium lacking histidine in the BRC-1^triA^ mutant, suggesting that while BRC-1^triA^ interacts with BRD-1, there is some impairment (Fig 4E). Quantitative analysis of an independent reporter, β-galactosidase, revealed a ∼50% decrease in the interaction between BRC-1^I23A^ and BRD-1 and a ∼90% interaction defect between BRC-1^triA^ and BRD-1 (Fig 4F). Thus, in contrast to human BRCA1, mutations in BRC-1 residues important for ligase activity also affect interaction with BRD-1 and this interaction defect may contribute to the more severe phenotype of *brc-1(triA)*.

### GFP fused to E3 ligase defective BRC-1 restores nuclear localization and partially rescues defects in DNA damage repair

In the course of our experiments we discovered that fusion of GFP to the N-terminus of the E3 ligase impaired *brc-1* mutants had less severe phenotypes in response to DNA damage compared to the non-tagged alleles, although only *gfp::brc-1(triA)* vs. *brc-1(triA)* reached statistical significance and was investigated further (p<0.0001; Fig 5A). Rescue in viability was specific to BRC-1, as C-terminal fusion of GFP to BRD-1 did not rescue embryonic lethality following IR in the E3 ligase defective mutants (S2B Fig). GFP::BRC-1 rescue was not a consequence of a change in BRC-1 expression, as there was no difference in steady state protein levels (S2C Fig). Interestingly, localization by live cell imaging revealed that unlike BRD-1 or BRD-1::GFP in the *brc-1(triA)* mutant, GFP::BRC-1^triA^ was enriched in the nucleus (-IR; Fig 5B, C), consistent with nuclear localization being important for function. As BRC-1-BRD-1 becomes enriched in nuclear foci in response to DNA damage (34, 35) and we observed improvement of function when GFP was fused to the BRC-1^triA^ following IR exposure, we examined localization of the mutant complex in response to IR-induced DNA damage. Two hours after exposure to IR, GFP::BRC-1^triA^ was largely nucleoplasmic as well as concentrated into nuclear foci in mitotic germ cells, although foci formation was not as robust as in wild type (+IR; Fig 5B). Interestingly, within the same germ line, no distinct GFP::BRC-1^triA^ foci were detected in meiotic nuclei following IR treatment. To quantify this, we calculated the coefficient of variation (CV), which provides a measure of the extent of foci above the nucleoplasmic GFP fluorescence signal. In mitotically-dividing *gfp::brc-1(triA)* mutant germ cells there was a significantly higher CV in IR treated worms compared to untreated worms; however, in meiotic cells there was no change in CV value following IR treatment (Fig 5D). These results suggest that independently recruiting BRC-1 to the nucleus by GFP fusion, where it can concentrate at DNA damage sites, partially suppresses the requirement for E3 ligase activity in proliferating germ cells.

**Fig 5.**
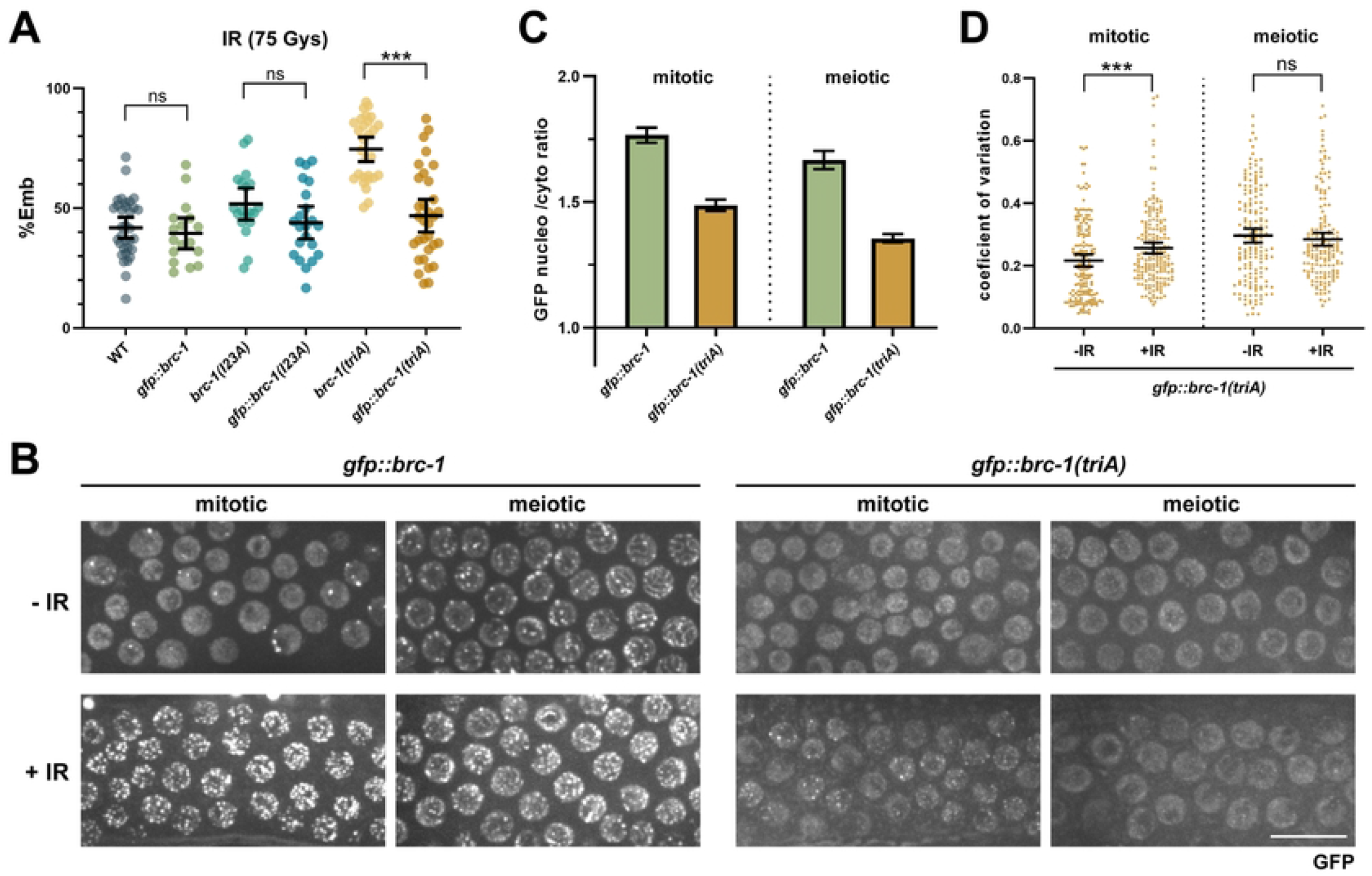
Nuclear accumulation of BRC-1 with impaired E3 ligase activity promotes viability in response to DNA damage. (A) Embryonic lethality in the presence of 75Gys IR was examined in wild type (n=33), *gfp::brc-1* (n=17), *brc-1(I23A)* (n=19), *gfp::brc-1(I23A)* (n=22), *brc-1(triA)* (n=28), and *gfp::brc-1(triA)* (n=32) animals. *** p < 0.001 Mann-Whitney. (B) Images of mitotic and meiotic (early pachytene -mid pachytene) germ cells expressing GFP::BRC-1 or GFP::BRC-1^triA^ in the absence (-IR) and presence (+IR) of 75Gys radiation. Scale bar=10μm. (C) Graph shows nucleoplasmic to cytoplasmic ratio of GFP signal in *gfp::brc-1* and *gfp::brc-1(triA)* strains. A minimum of 60 nuclei from 3 germ lines were analyzed. (D) Coefficient of variation for GFP::BRC-1^triA^ fluorescence to reflect changes in localization (foci formation) in response to IR in mitotic and meiotic nuclei in the germ line; five germ lines were analyzed for each genotype. Statistical comparisons between - and + IR *** p < 0.001 Mann-Whitney.

### BRC-1-BRD-1 E3 ligase activity is essential for recruitment of the complex to meiotic DSBs

In contrast to mitotic germ cells, essentially no GFP::BRC-1^triA^ foci were observed in meiotic nuclei after exposure to IR, suggesting that meiosis is more sensitive to loss of E3 ligase activity in recruiting the complex to DNA damage sites (Fig 5B, D). To probe the requirement for BRC-1-BRD-1 E3 ligase activity in recruitment of the complex to meiotic DSBs, we monitored GFP::BRC-1^triA^ localization in the *syp-1* mutant, where homologous chromosomes fail to synapse and no crossovers are formed (48). As we previously reported, there were extensive GFP::BRC-1 nuclear foci in the *syp-1* mutant; these foci presumably represent meiotic recombination sites that are delayed in repair due to the absence of a homologous repair template (34). In contrast to GFP::BRC-1, essentially no GFP::BRC-1^triA^ foci were observed in the *syp-1* mutant (Fig 6A, D). This result suggests that E3 ligase activity is critical for accumulation of BRC-1-BRD-1 at sites of meiotic recombination when chromosomes fail to synapse.

**Fig 6.**
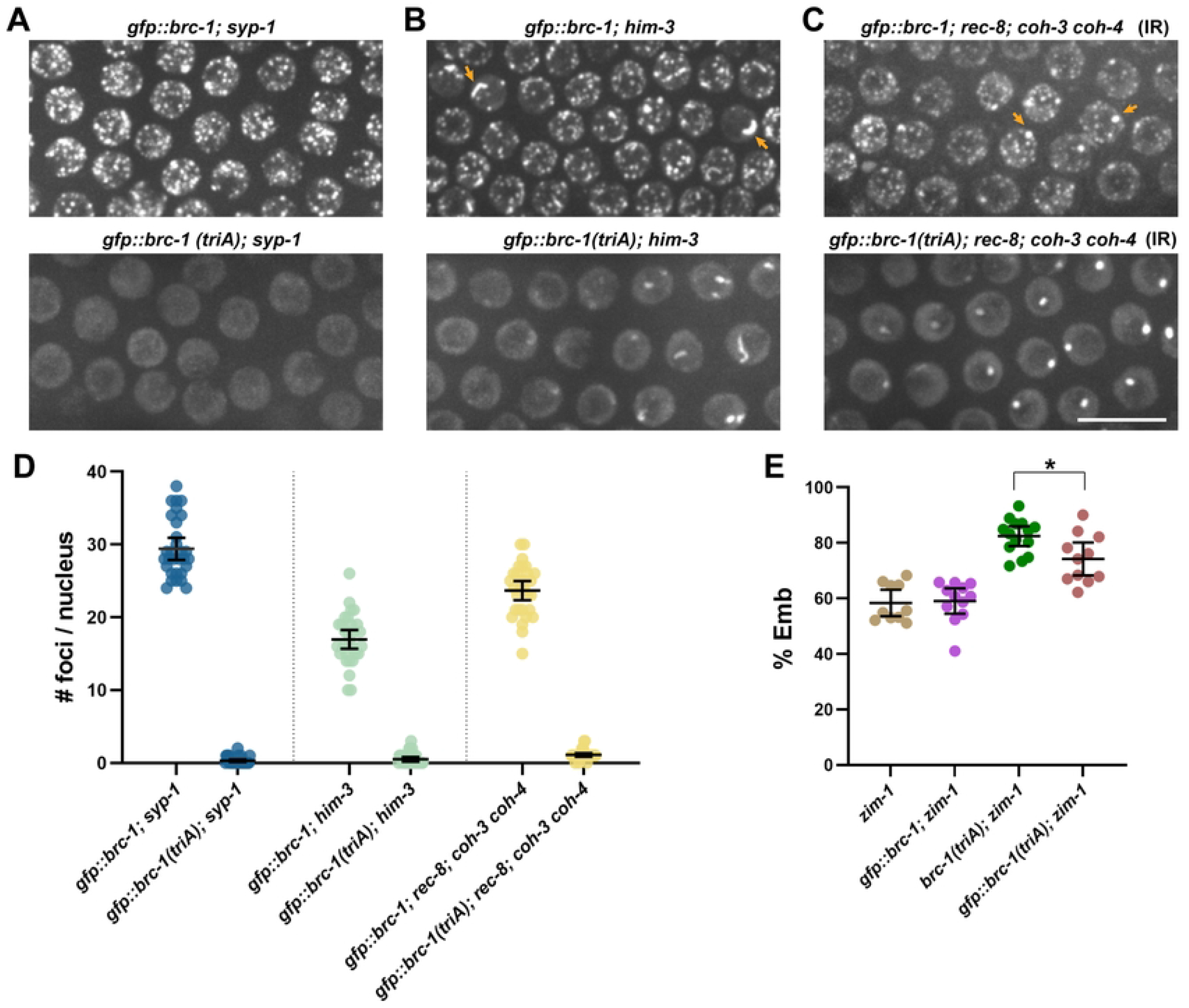
BRC-1-BRD-1 E3 ligase activity is essential for recruitment of the complex to DSBs on meiotic chromosomes. Images of meiotic (early pachytene - mid pachytene) germ cells expressing GFP::BRC-1 or GFP::BRC-1^triA^ in *syp-1* (A) and *him-3* (B) mutants without IR, and *rec-8; coh-3 coh-4* mutants in the presence of 75Gys IR (C). Scale bar=10μm. (D) Quantification of GFP::BRC-1 or GFP::BRC-1^triA^ foci observed in the different mutants; a minimum of 3 germ lines from half-projections were scored. (E) Embryonic lethality in *zim-1* (n=10), *gfp::brc-1; zim-1* (n=12), *brc-1(triA); zim-1* (n=14), *gfp::brc-1(triA); zim-1* (n=11) mutants. * p < 0.05 Mann-Whitney.

To determine whether the more severe defect in recruitment of GFP::BRC-1^triA^ to sites of recombination in meiosis was a consequence of barriers imposed by the specialized meiotic chromosome structure, we examined the requirement for BRC-1-BRD-1 E3 ligase activity in recruitment of the complex to DSBs in mutants defective in the formation of the chromosome axes. To that end, we monitored the localization of GFP::BRC-1 and GFP::BRC-1^triA^ when axis formation was impaired by mutation of the HORMA domain protein, HIM-3. HIM-3 is an axis component and is required for homolog pairing and synapsis and promotes crossover formation by biasing recombination to the homologous chromosome instead of the sister chromatid (50, 51). While GFP::BRC-1 is recruited to both foci and the occasional track in the *him-3* mutant, as we have observed previously in mutants defective in meiotic recombination (e.g., *spo-11, mre-11, msh-5*) (33, 34), only tracks but no GFP::BRC-1^triA^ foci were observed in *him-3* (Fig 6B, D). Quantification of GFP::BRC-1 foci showed that the average number was lower in the *him-3* mutant compared to *syp-1* (29.4 ± 4.0 vs. 16.9 ± 3.5; p<0.0001), consistent with repair being more efficient in *him-3* mutants due to release of the barrier to inter-sister repair (Fig 6D). We also examined the consequence of impairing meiotic chromosome cohesion and hence axis formation by mutation of the meiosis-specific cohesin kleisin subunits, REC-8, COH-3, and COH-4 (52, 53). Unlike *syp-1* and *him-3* mutants, *rec-8; coh-3 coh-4* triple mutants are not competent for meiotic DSB formation and therefore no GFP::BRC-1 foci were detected (53).

Consequently, we monitored recruitment of GFP::BRC-1 and GFP::BRC-1^triA^ to DNA breaks induced by IR and while we observed abundant GFP::BRC-1 foci, no GFP::BRC-1^triA^ foci were detected (Fig 6C, D). We also observed a bright aggregate of both GFP::BRC-1 and GFP::BRC-1^triA^ in the *rec-8; coh-3 coh-4* mutant in the presence and absence of IR, which is likely SC-like structures formed independently of chromosomes (polycomplexes). These results suggest that the chromosome axis does not impose a special requirement for BRC-1-BRD-1 E3 ligase-dependent recruitment of the complex to DSBs.

We next examined the phenotypic consequence of the inability to recruit nuclear GFP::BRC-1^triA^ to meiotic foci by examining progeny viability in the *zim-1* mutant. We observed improved progeny viability of GFP::BRC-1^triA^ compared to BRC-1^triA^, but not to the extent of what was observed in response to IR (Fig 6E and Fig 5A). Thus, nuclear BRC-1-BRD-1 provides some function despite its inability to accumulate at DSBs. Taken together, meiosis imposes a special requirement for BRC-1-BRD-1 E3 ligase activity for recruitment to DNA damage and meiotic recombination sites.

### BRD-1 function can be partially bypassed by expressing GFP::BRC-1

The partial rescue of the E3 ligase impaired mutants by fusing GFP to BRC-1, but not to BRD-1, prompted us to explore the contribution of BRD-1 to the function of the complex. To that end, we constructed a null allele (*brd-1(null)*) by engineering multiple stop codons in the second exon of *brd-1* as described (54), as available alleles of *brd-1* are in-frame deletions C-terminal to the RING domain and helices where the two proteins interact (34). *brd-1(null)* mRNA was unstable and no GFP fluorescence was detected in *brd-1(null)* worms containing GFP fused to the C-terminus of *brd-1*, providing evidence that it is a null allele (S3A, B Fig). Further, *brd-1(null)* was phenotypically indistinguishable from *brc-1(null)* for male self-progeny and embryonic lethality in the absence and presence of IR (S3C and S4A Figs).

We next examined the phenotype of *gfp::brc-1 brd-1(null)* and saw a partial rescue of embryonic lethality following exposure to IR (Fig 7A). Consistent with this, GFP::BRC-1 was enriched in the nucleus and formed weak foci in response to IR in the absence of BRD-1, although there was reduced steady state levels of GFP::BRC-1 compared to in the presence of BRD-1 (Fig 7B, C; S4 Fig). These findings suggest that GFP::BRC-1 alone can provide some function without its binding partner. Rescue was specific to appending GFP to BRC-1, as neither GFP::BRD-1 nor BRD-1::GFP could provide partial function when BRC-1 was absent as measured by embryonic lethality following IR treatment (S4A Fig). Additionally, neither GFP::BRD-1 nor BRD-1::GFP accumulated in the nucleus in the absence of BRC-1 (34) (S4B Fig).

**Fig 7.**
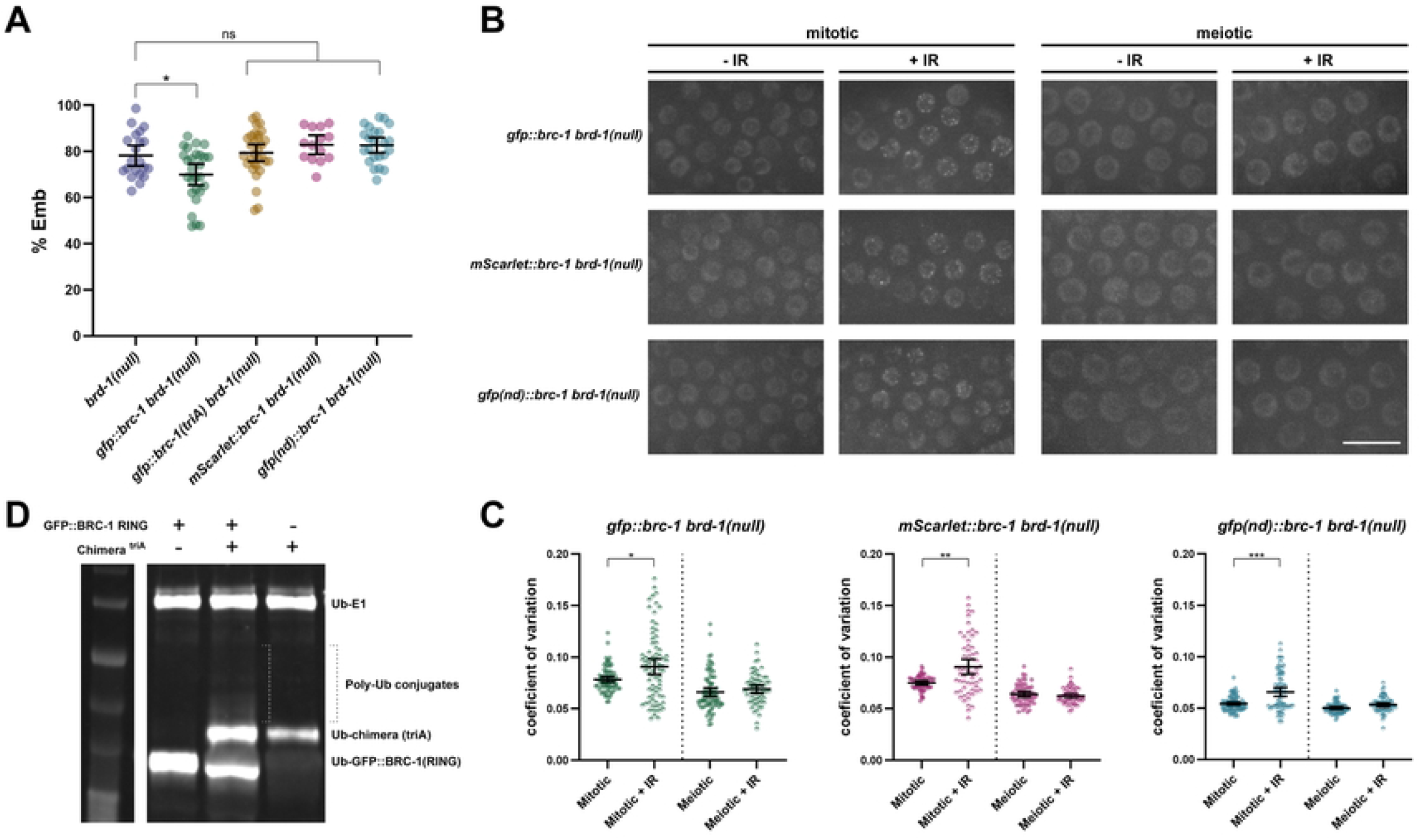
BRC-1 nuclear accumulation and self association are required to partially bypass the requirement for BRD-1. (A) Embryonic lethality in the presence of 75Gys IR was examined in *brd-1(null)* (n=21), *gfp::brc-1 brd-1(null)* (n=27), *gfp::brc-1(triA) brd-1(null)* (n=31), *mScarlet::brc-1 brd-1(null)* (n=14), *gfp(nd) brd-1(null)* (n=23). * p < 0.05 by Mann-Whitney. (B) Images of mitotic and meiotic (early pachytene - mid pachytene) *brd-1(null)* germ cells in the absence (-IR) and presence of IR (+IR) expressing either GFP::BRC-1, mScarlet::BRC-1 or GFP^nd^::BRC-1. Scale bar = 10μm. (C) Coefficient of variation for GFP::BRC-1, mScarlet::BRC-1, and GFP^nd^::BRC-1 fluorescence to reflect changes in localization (foci formation) in response to IR in the absence of BRD-1 in mitotic and meiotic germ cell nuclei; a minimum of 4 germ lines were analyzed for each genotype. Statistical comparisons between - and + IR by Mann-Whitney: * p < 0.05, ** p < 0.01, *** p < 0.001 by Mann-Whitney. (D) Immunoblot of auto-ubiquitylation (anti-HA-Ub) of GFP::BRC-1 RING, GFP::BRC-1 RING in the presence of the BRD-1-BRC-1^triA^ chimera, and BRD-1-BRC-1^triA^ chimera alone. ∼2x more monoubiquitylation on the BRD-1-BRC-1^triA^ chimera as well as polyubiquitylation were observed when GFP::BRC-1 RING was included in the reaction.

To determine the consequence of removing BRD-1 to BRC-1 E3 ligase activity we expressed and purified GFP::BRC-1 RING in *E. coli* (S4C Fig) and assayed auto-ubiquitylation *in vitro*. We observed significant auto-monoubiquitylation of GFP::BRC-1 RING, but no polyubiquitylation, suggesting that GFP::BRC-1 RING alone is competent to transfer a single ubiquitin onto itself (Fig 7D). To ascertain whether the lack of polyubiquitylation was due to the absence of BRD-1, we performed the *in vitro* assay using GFP::BRC-1 RING in the presence of the BRD-1-BRC-1^triA^ chimera, which is incapable of polyubiquitylation (Fig 3B, C). Addition of BRD-1-BRC-1^triA^ to the reaction resulted in both an increase in monoubiquitylation of the chimera and some polyubiquitylation (total ubiquitin signal was 2x the BRD-1-BRC-1^triA^ ubiquitin signal alone; Fig 7D). These results suggest that BRC-1 can monoubiquitylate itself when fused to GFP in the absence of BRD-1, but that BRD-1 is required for polyubiquitylation of the complex.

To determine whether GFP::BRC-1-mediated monoubiquitylation was functionally important, we expressed GFP::BRC-1^triA^ in *brd-1(null)* and monitored embryonic lethality following exposure to IR. No significant rescue was observed, suggesting that BRC-1-mediated monoubiquitylation is important in response to DNA damage (Fig 7A).

### BRC-1 nuclear accumulation and self-association are important for function in the absence of BRD-1

To ascertain how fusion of GFP to the N-terminus of BRC-1 promotes function in the absence of BRD-1, we constructed an N-terminal fusion with mScarlet, a monomeric red fluorescent protein (55), at the endogenous BRC-1 locus. The mScarlet::BRC-1 fusion was fully functional (S4A Fig). However, expression of mScarlet::BRC-1 in *brd-1(null)* did not improve progeny viability following exposure to IR, even though mScarlet::BRC-1 was nuclear, could form foci in response to IR, and was as stable as GFP::BRC-1 in the *brd-1(null)* mutant (Fig 7B, C, S4D Fig). These results suggest that nuclear accumulation, while necessary, is not sufficient for BRC-1 function independent of BRD-1. As the name implies, mScarlet is monomeric, while GFP has the tendency to dimerize or oligomerize, particularly at high concentrations (56). We next addressed whether association between GFP molecules was important for bypassing BRD-1 function. To that end, we modified the GFP fused to BRC-1 by substituting hydrophobic amino acids with charged amino acids on the surface interface (A206K, L221K, F223R) (57); we refer to this as GFP^nd^::BRC-1 (nd for non-dimerizable). As with mScarlet::BRC-1, GFP^nd^::BRC-1 is fully functional in an otherwise wild-type worm (S4A Fig). However, expression of GFP^nd^::BRC-1 did not provide any rescue of the *brd-1(null)* mutant even though it was nuclear, could form foci in response to IR, and was as stable as GFP::BRC-1 in the *brd-1(null)* mutant (Fig 7A, B, C; S4D Fig). These results indicate that nuclear accumulation and self-association of BRC-1 driven by GFP can partially bypass the requirement for BRD-1 in response to DNA damage.

## Discussion

Here we take advantage of *C. elegans* to examine the requirement for BRCA1-BARD1 E3 ubiquitin ligase activity *in vivo* in the context of DNA damage signaling and meiosis. We find that mutants significantly impaired for E3 ligase activity *in vitro* still provide some function *in vivo*. We provide evidence that nuclear localization and BRC-1-BRD-1 association are critical for the function of the complex and these properties are impacted when E3 ligase activity is abrogated. Additionally, we show that GFP fusion to BRC-1 can drive protein accumulation in the nucleus and BRC-1 self-association, which partially rescue defects in DNA damage repair in the absence of BRD-1, indicating that BRC-1 is the key functional unit of the complex, while BRD-1 plays an accessory role to augment BRC-1 function.

### A BRCA1 ligase dead mutant?

The role of BRCA1-BARD1 E3 ubiquitin ligase activity has remained enigmatic, due in part to the absence of a true ligase dead allele (27-29). Based on extensive biochemical and structural work on RING-type E3 ligases in general, and the human BRCA1-BARD1 complex specifically, we constructed two BRC-1 mutants predicted to interfere with E3 ligase activity: I23A and triA (14, 23, 29, 58, 59) (Fig 1A, B). In human BRCA1 isoleucine 26 defines the binding site for E2 conjugating enzymes, while lysine 65 is the linchpin residue that activates E2-ubiquitin for ubiquitin transfer; human BRCA1 harboring the triple I26A, L63A, K65A mutation is an E3 ligase dead mutant *in vitro* (29). The corresponding isoleucine 23 and arginine 61 residues in *C. elegans* BRC-1 likely play analogous roles in E2 binding and activation of E2-ubiquitin and therefore the triA mutant is predicted to be a ligase dead enzyme (Fig 1B). Surprisingly, while both BRC-1^I23A^ and BRC-1^triA^ are significantly impaired for E3 ligase activity *in vitro* (Fig 3), they have different phenotypes *in vivo* (Figs 1, 2, 4). Further, neither *brc-1(I23A)* nor *brc-1(triA)* has a phenotype as severe as *brc-1(null)*, suggesting that in addition to E3 ligase activity, the complex serves a structural role to promote DNA damage signaling, repair, and meiotic recombination. This is consistent with studies of human BRCA1, where RING-less mutants maintain some homologous recombination function (60, 61).

Human BRCA1-BARD1 is capable of coupling with multiple E2s *in vitro* and different E2s define mono vs. polyubiquitylation of substrates and how polyubiquitin chains are linked to each other. The BRCA1^I26A^ mutant has residual E3 ligase activity with a subset of E2s, including UbcH5c, which in complex with BRCA1-BARD1 promotes polyubiquitylation (23, 29). We used UbcH5c in *in vitro* ubiquitylation assays and observed robust auto-polyubiquitylation as well as ubiquitylation of H2A with the wild-type chimera but no detectable self-polyubiquitylation with either I23A or triA chimeras, nor any ubiquitylation of histone H2A (Fig 3). However, there was significantly reduced but detectable auto-monoubiquitylation with the mutant chimeras. It is not clear whether monoubiquitylation represents residual activity that is only modestly reduced by mutation of I59A and R61A in the I23A mutant in *C. elegans* BRC-1 or is perhaps a consequence of the RING domains of BRD-1 and BRC-1 being physically tethered in the chimera. We did observe low levels of monoubiquitylation in the absence of any E2 (S1 Fig), suggesting that some auto-monoubiquitylation may result from enhanced interaction between BRC-1 and BRD-1 RING domains within the chimera.

UbcH5c is orthologous to *C. elegans* UBC-2 (LET-70), which has previously been shown to couple with BRC-1-BRD-1 for ubiquitin transfer in the context of DNA damage signaling (35). Similar to human BRCA1-BARD1, it is likely that *C. elegans* BRC-1-BRD-1 couples with multiple E2s to regulate different pathways (e.g., DNA damage signaling, meiosis, heterochromatin regulation, axon outgrowth) (31-37, 41, 42). The *C. elegans* genome encodes 22 E2s and the entire spectrum of these E2s coupling to different E3 ligases is not clear (62). Recently developed tools to conditionally deplete proteins in a tissue-specific manner could help define how different E2s couple with BRC-1-BRD-1, and other E3 ubiquitin ligases, to regulate different pathways *in vivo* (63-65).

While *C. elegans* BRC-1-BRD-1 shares many similarities with the human complex, it is not surprising that differences have evolved between worms and humans. For example, we found that in contrast to the human proteins, *C. elegans* E3 ligase defective BRC-1 show impaired interaction with BRD-1 (29) (Fig 4). One possibility is that these amino acid substitutions directly alter how BRC-1 and BRD-1 interact, although these do not reside in the helices required for binding between BRC-1 and BRD-1. Alternatively, BRC-1-BRD-1 auto-ubiquitylation may enhance interaction between these proteins, consequently ubiquitylation and interaction are impaired in E3 ligase defective BRC-1. Using physically tethered BRC-1 and BRD-1 RING domains in our chimeric proteins in the *in vitro* assay may have masked the interaction defect, leading to similar impairment of E3 ligase activity *in vitro* in BRC-1^I23A^ and BRC-1^triA^, but different phenotypes *in vivo*. Nonetheless, continued analyses in *C. elegans* will be instrumental in defining the fundamental roles of BRC-1-BRD-1 in the context of a whole organism.

### BARD1 serves an accessory role to ensure BRCA1-mediated polyubiquitylation and nuclear localization

BARD1 was identified as a BRCA1 interacting protein and mutations in BARD1 also lead to an increased incidence of cancer (2, 6, 66). Structural work defined the contact sites between the two proteins at the helices adjacent to the RING domains and demonstrated that only BRCA1 binds E2s for ubiquitin transfer, while BARD1 is required to stimulate BRCA1’s E3 ligase activity (15, 67, 68). We observed robust auto-polyubiquitylation of the wild-type chimera *in vitro;* however, assaying GFP::BRC-1 RING alone revealed significant auto-monoubiquitylation only, but no polyubiquitylation, in the presence of the same E2. Addition of the triA chimera to the GFP::BRC-1 RING reaction promoted the formation of polyubiquitylation, suggesting that BRD-1 is specifically required for BRC-1-mediated polyubiquitylation. Whether BARD1 also stimulates BRCA1 polyubiquitylation in mammalian cells is unclear; however, it has been shown that BRCA1-BARD1 auto-polyubiquitylation enhances the E3 ligase activity of the full-length complex *in vitro* (14, 15, 69).

In addition to promoting E3 ligase activity, BARD1 is important for the stability and nuclear retention of BRCA1 *in vivo*. Analysis of human BRCA1-BARD1 have revealed multiple mechanisms, including both regulated nuclear import and export driven by interaction between the two proteins, to ensure nuclear localization of the complex where it primarily functions (70, 71). Similar to what has been reported in mammals, *C. elegans* BRD-1 is required for the stability and nuclear localization of BRC-1 (32, 72). It was therefore surprising that appending GFP to the N-terminus of BRC-1 could bypass the requirement for BRD-1 in promoting nuclear accumulation of BRC-1. GFP::BRC-1 alone could partially promote DNA damage signaling in the absence of BRD-1 and this was dependent on both nuclear localization and self-association driven by GFP. These results reinforce that BRCA1 is the primary functional unit of the complex and its key functions and targets are within the nucleus, while BARD1 serves an accessory role to bolster BRCA1-mediated polyubiquitylation and nuclear localization.

While both BRCA1 and BARD1 possess N-terminal RING and C-terminal BRCT domains, BARD1 uniquely contains conserved ankyrin repeats in the middle of the protein (10). Recent molecular and structural studies have revealed that the BARD1 ankyrin and BRCT domains direct the interaction of BRCA1-BARD1 to N-terminal ubiquitylated histone H2A within the nucleosome, a chromatin mark associated with DSBs. Once bound, the complex mediates the ubiquitylation of the C-terminal tail of H2A, which opposes the binding of 53BP1 to promote repair by homologous recombination (20-22). Given the unique requirement for BARD1 ankyrin domains in recruitment to damaged DNA, how does GFP::BRC-1 partially bypass the need for BRD-1 with respect to DNA damage signaling? One possibility is that there are redundant mechanisms for recruitment of BRC-1-BRD-1 to DSBs. Human BRCA1-BARD1 recruitment to DNA damage sites has been shown to be mediated through both BRCA1-BARD1 and RAP80 (73). While no obvious RAP80 ortholog has been identified in *C. elegans*, other interacting proteins may serve a similar role in the recruitment of the complex to DSBs, and/or sequences within BRC-1 itself may facilitate concentration at DNA damage sites.

### BRC-1-BRD-1 E3 ligase activity is required for recruitment of the complex to meiotic DSBs

Our analysis of the E3 ligase defective mutants revealed that while GFP fusion to BRC-1^triA^ drives nuclear accumulation and the protein is capable of foci formation in response to DNA damage in the mitotic germ cells, E3 ligase activity is critical for the recruitment of the complex to DSBs in meiotic cells. Unique to meiosis is the pairing and synapsis of homologous chromosomes, which are essential for crossovers formation to ensure that the homologs segregate properly at Meiosis I. These events occur within the specialized structure of meiotic chromosomes, which includes the chromosome axes and the SC. Chromosome axes are extended filaments, which provide a scaffold for the organization of chromosomes as a linear array of loops (74, 75), and become the lateral elements of the SC. We found that blocking the formation of the chromosome axis, or the SC, did not alleviate the requirement for BRC-1-BRD-1 E3 ligase activity, suggesting that their presence does not impose an additional barrier for recruitment of the complex to meiotic DSBs (Fig 6). In addition to the specialized structure of meiotic chromosomes, the chromatin landscape is also different in meiotic cells and this unique chromatin environment may dictate the requirement for E3 ligase activity in recruiting the complex to meiotic DSBs (76). Additionally, context-specific BRC-1-BRD-1 post-translational modifications and/or interacting proteins may exist that define redundant pathways for recruiting the complex to DNA damage sites in mitotic germ cells. Future work will provide insight into the context-dependent recruitment of BRC-1-BRD-1 in response to DNA damage.

## Conclusion

BRCA1-BARD1 regulates a plethora of processes *in vivo* and mounting evidence indicates that BRCA1-BARD1 E3 ligase activity is critical for several aspects of the complex’s function, including tumor suppression. However, the underlying molecular mechanisms are just beginning to be revealed. Our findings that BRC-1 is the key driver for DNA damage signaling and repair within the heterodimer is consistent with the observed higher prevalence of pathogenic variants identified in BRCA1 as compared to BARD1 (77, 78). Further, mutations in BRCA1 pose high risk for both breast and ovarian cancer, while BARD1 mutations are only a risk factor for breast, but not ovarian cancer (79-81). Thus, as in *C. elegans*, human BRCA1 and BARD1 are not equivalent in function leading to different spectrum of cancers when mutated.

## Materials and methods

### Genetics

*C. elegans* strains used in this study are listed in Supplemental Table 1. Some nematode strains were provided by the Caenorhabditis Genetics Center, which is funded by the National Institutes of Health National Center for Research Resources (NIH NCRR). Strains were maintained at 20°C.

### CRISPR-mediated allele construction

*brc-1(xoe4), gfp::brc-1(xoe7)* and *brd-1::gfp(xoe14)* have been described (34). *gfp::brc-1(xoe20[I23A])* and *mScarlet-i::brc-1(xoe34)* were generated using CRISPR-mediated genome editing with a self-excising cassette as described in (82) with modifications as follows: I23A was introduced at the same time with GFP knock-in by incorporating the corresponding mutation in the 3’ homology arm on the repair template plasmid using the Q5 site-directed mutagenesis kit (New England Biolabs). <u>G</u>erm<u>L</u>ine <u>O</u>ptimized mScarlet-i sequence (Fielmich *et al*. 2018) was cloned into the repair template plasmid in place of GFP by Gibson Assembly to generate *mScarlet-i::brc-1(xoe34). gfp::brc-1(xoe48[triA]*) was generated by introducing the corresponding I59A R61A mutations in the *gfp::brc-1(xoe20[I23A])* background using the co-CRISPR method (83). All other genome-edited strains were generated using the co-CRISPR method. *brd-1(xoe58[null-gfp::3xFLAG])* was generated by introducing the stop-in cassette into *brd-1::gfp(xoe14)* (34). Guide sequence, repair template, and primers for genotyping are provided in Supplemental Table 2. All strains were outcrossed for a minimum of three times before analyses.

### Embryonic lethality in the absence and presence of irradiation and male self-progeny

L4 hermaphrodites were transferred to individual plates (-IR) or exposed to 75Gys γ-irradiation from a ^137^Cs source, and then transferred to individual plates. Individually plated hermaphrodites were transferred to new plates every 24hr for 3 days. Embryonic lethality was determined by counting eggs and hatched larvae 24hr after removing the hermaphrodite and calculating percent as eggs/(eggs + larvae). Males were scored after 72hr and calculating percent as males/(males + hermaphrodites + eggs).

### Cytological analyses

#### Immunolabeling

Germ lines were immunolabeled as described (84). The following primary antibodies were used at the indicated dilutions: rabbit anti-RAD-51 (2948.00.02; SDIX; 1:5,000; RRID: AB_2616441), rabbit anti-BRD-1 (1:500; from Dr. Simon Boulton(35)). Secondary antibodies Alexa Fluor 594 donkey anti-rabbit IgG from Life Technologies were used at 1:500 dilution. DAPI (2μg/ml; Sigma-Aldrich) was used to counterstain DNA.

#### Image capture

Collection of fixed images was performed using an API Delta Vision Ultra deconvolution microscope equipped with an 60x, NA 1.49 objective lens, and appropriate filters for epi-fluorescence. Z stacks (0.2μm) were collected from the entire gonad. Images were deconvolved using Applied Precision SoftWoRx batch deconvolution software and subsequently processed and analyzed using Fiji (ImageJ) (Wayne Rasband, NIH).

For live cell imaging, 18–24 hr post L4 hermaphrodites were anesthetized in 1mM tetramisole and immobilized between a coverslip and a 2% agarose pad on a glass slide. Z-stacks (0.33μm) were captured on a spinning-disk module of an inverted objective fluorescence microscope with a ∼100Å, NA 1.46 objective, and EMCCD camera. Z-projections of stacks were generated, cropped, and adjusted for brightness in Fiji.

#### RAD-51 foci quantification

RAD-51 foci were quantified in a minimum of three germ lines of age-matched hermaphrodites (18-24hr post-L4). As the *zim-1* mutation results in an extended transition zone, we divided germ lines into four equal zones beginning from the first row with two or more crescent-shaped nuclei until the end of diplotene (Fig 2B). RAD-51 foci were quantified from half projections of the germ lines; the number of foci per nucleus was scored for each zone. To measure pixel intensities of RAD-51, foci were identified by a prominence value between 10-20 using the “Find Maxima” function embedded in Fiji from half projections of germ lines. Pixel intensities were measured using ROI Manager in Fiji from defined regions of the gonad and the values were plotted on scatterplot with means and 95% CI using GraphPad Prism. A minimum of three germ lines for each genotype were used for quantification.

#### Nuclear to cytoplasmic ratio

Mean pixel intensity of BRD-1 immunolabeling or direct GFP fluorescence was measured from 80 nuclei and surrounding cytoplasm from three different germ lines in mitotic and meiotic (early to mid pachytene) regions of the gonad using Fiji. The nucleoplasmic to cytoplasmic ratio was calculated and the mean and 95% CI for the ratio was plotted.

#### Coefficient of variation

GFP::3xFLAG::BRC-1^triA^ fluorescence following exposure to 75Gys IR was quantified by measuring the mean fluorescence intensity and standard deviation (SD) in Fiji for individual nuclei [region of interest (ROI)] in mitotic germ cells (proliferative zone) and meiotic germ cells (early to mid pachytene). Coefficient of variation (CV) is defined as SD of intensity divided by mean intensity (85). The CV describes the dispersion of pixel intensity values from a 2D ROI around the mean pixel intensity such that nuclei with more distinct foci will have high CV values, whereas nuclei with more uniform fluorescence will have low CV values.

#### GFP::BRC-1, and GFP::BRC-1^triA^ foci quantification

Foci were quantified in 10 mid pachytene nuclei from each of three half projections of germ lines of age-matched hermaphrodites (18-24hr post-L4).

#### Protein Constructs

The BRD-1-BRC-1 chimera, encoding amino acids 1-107 of BRD-1 and amino acids 2-106 of BRC-1 separated by a GGSGG-linker was synthesized as a G-block and cloned into pET28A vector containing a PreScission protease cleavage site, superfolder GFP (sfGFP) and a strepII-tag, using Gibson Assembly. Mutant BRD-1-BRC-1 chimeras harboring the single I23A and triA mutations were similarly synthesized as G-blocks and cloned into pET28A as described above. The GFP::BRC-1 RING sequence in pET28A encodes amino acids 2-106 of BRC-1 and a N-terminal GFP followed by 3x FLAG-tag and a C-terminal strepII-tag. The protein expressed from this construct has identical amino acid sequences as the fusion protein expressed in *gfp::brc-1(xoe7)* allele with the exception of truncated BRC-1 RING domain and the addition of the strepII-tag.

#### Protein purification

The wild-type and mutant BRD-1-BRC-1 chimeras were expressed in BL21-CodonPlus (DE3)-RIPL cells (Agilent). The cells were grown at 37°C until OD^600^ 0.6 and were induced by 0.2mM isopropyl-β-D-thiogalactoside in the presence of 100μM ZnCl_2_ at 37°C for 6 hrs. The GFP::BRC-1 RING was induced overnight at 18°C. After induction, cells were harvested and resuspended in buffer A (20mM HEPES-KOH pH 7.2, 300mM NaCl, 1mM EGTA) supplemented with 1mM DTT, 0.2% NP-40, protease inhibitors (1mM PMSF; protease inhibitor cocktail P83340; Sigma-Aldrich) and lysed using a Emulsiflex C-3 (Avestin) high pressure homogenizer. The lysates were centrifuged at 15000xg for 20min at 4°C. The supernatants were passed through Strep-Tactin XT (IBA) for affinity purification, and the column was washed with lysis buffer to remove unbound proteins before eluting the proteins with 50 μM biotin (Chem-Impex Int’l) in low salt buffer (20mM HEPES-KOH pH 7.2, 30 mM NaCl, 0.1% NP40). Proteins were further purified by anion exchange using HiTrap Q HP column equilibrated with 20mM HEPES-KOH pH7.2 with a linear NaCl gradient from 0mM to 600mM. Peak fractions were pooled and concentrated on Amicon-Ultra spin filters (EMD Millipore) and supplemented with 10% glycerol. Protein aliquots were snap frozen in liquid nitrogen and stored at −80°C. Protein concentration was measured using a Nanodrop One (ThermoFisher) based on the total amount of fluorophore (sfGFP or GFP).

#### E3 ligase activity assay

Ubiquitin transfer reactions were performed in 30μl reaction volume at 30°C for the indicated time with mild rocking. For end point auto-ubiquitylation assays, the reaction mixture contained 0.2μM E1 (hUBE1; E-305; bio-techne), 1μM E2 (hUbcH5c; E2-627; biotechne), 5μM BRD-1-BRC-1 chimera, 20μM HA-ubiquitin (U-110; bio-techne), 5mM ATP, 5mM MgCl_2_ in reaction buffer (20mM Hepes pH 7.2; 150mM NaCl). To test ubiquitylation of histone H2A, 0.75μM human histone H2A (ab200295; Abcam) was added to the above reaction mixture. For time course experiments, 0.1μM E1, 0.5μM E2, 3μM BRD-1-BRC-1 chimera, 10μM HA-ubiquitin, 5mM ATP, 5mM MgCl_2_ were mixed in a 150μl reaction volume and incubated with mild rocking at 30°C. 30μl were removed at 0, 5, 10, 20, 40min and the reactions stopped with 10μl 4X sample buffer followed by boiling. Reaction mixtures were visualized by immunoblot and analyzed by measuring pixel intensity of ubiquitylated species.

#### Immunoblot analysis

For steady state protein levels, whole worm lysates were generated from indicated genotypes. ∼200 worms were collected in M9 buffer, washed 2x in M9 and then resuspended in equal volume of 2X Laemmli sample buffer (Bio-RAD) in a total volume of 40μl. Worm lysates or E3 ligase reaction mixtures were resolved on 4-20% stain-free SDS-PAGE gels (Bio-RAD) and transferred to Millipore Immobilon-P PVDF membranes. Membranes were blocked with 5% nonfat milk and probed with mouse anti-FLAG (MA1-91878; Invitrogen; 1:1000; RRID AB_1957945), rabbit anti-GFP (NB600-308; Novus Biologicals; 1:2000; RRID: AB_10003058), mouse anti-HA [12CA5; amino acids 98–106 of human influenza virus hemagglutinin protein; IgG2b mAb; 1:1000; RRID: AB_2532070; in-house (Trimmer Laboratory)], or rabbit anti-Histone-H2A (ab18255; Abcam; 1:1000; RRID:AB_470265) followed by IRDye800-conjugated anti-mouse IgG secondary antibodies (962 32212; LI-COR Bioscience; 1:20000; RRID: AB_621847) or IRDye680-conjugated anti-rabbit IgG secondary antibodies (925-68073:; LI-COR Bioscience; 1:20000; RRID: AB_2716687). Immunoblots were imaged on a LI-COR Odyssey Infrared Imager, signal was quantified using Image StudioLite and normalized with total protein input measured from the stain-free signal using BioRad Gel Doc™ EZ System.

#### Yeast two-hybrid

Full length wild-type or mutant BRC-1 sequences were cloned into plasmid pBridge (Takara Bio), transformed into yeast strain Y2HGold (Takara Bio) and transformants were selected on medium lacking tryptophan. Full length BRD-1 sequences were cloned into plasmid pACT2.2, transformed into yeast strain Y187 (Takara Bio) and transformants were selected on medium lacking leucine. Wild type or mutant BRC-1 expressing strains were mated with BRD-1 expressing strain and the diploids selected on -Trp - Leu double dropout plate at 30°C. Diploid cells were grown in liquid -Trp - Leu double dropout medium overnight, and serial dilutions were plated on -His -Trp -Leu triple dropout and -Trp -Leu double dropout solid media. For quantitative measurement of wild type or mutant BRC-1 and BRD-1 interactions, β-galactosidase activity was measured. Cell lysates were incubated in the presence of CPRG (chlorophenol red-β-D-galactopyranoside, RocheApplied Science Cat. NO.10884308001) as substrate, color change was measure at OD_578_ and β-galactosidase units were calculated as described (Yeast Protocol Handbook, Takara Bio).

#### RT-PCR

Total RNA was isolated from 50 to 100 μl of packed worms from wild type and *brd-1(null)* using the RNeasy Mini Kit (74104; Qiagen) and QIAshredder (79654; Qiagen). 1 μg of RNA was converted to cDNA using SuperScript III First-Strand Synthesis System for RT-PCR (18080-051; Invitrogen) primed with Oligo(dT)20. PCR was performed in a standard PCR machine with 20 cycles of amplification and resolved by gel electrophoresis.

## Acknowledgments

We thank the Caenorhabditis Genetic Center, which is funded by NIH Office of Research Infrastructure Programs (P40 OD010440) for providing strains. We thank Dr. Satoshi Namekawa (University of California Davis) for the histone H2A antibody and Brian Wong for constructing strains. We are particularly grateful to Dr. Judy Callis (University of California Davis) for input on E3 ligase activity assays, Dr. Daniel Elatan (University of California Davis) for help with AlphaFold and ChimeraX as well as imaging analyses, and the Engebrecht lab for thoughtful discussions. We thank the MCB Light Microscopy Imaging Facility, which is a UC Davis Campus Core Research Facility, for the use of the Deltavision Ultra and 3i Spinning Disc microscopes for generating images. This work was supported by National Institutes of Health GM103860 and GM103860S1 to JE and 2R35GM124889-06 to RJM.

## Fig Legends

**S1 Fig. BRD-1-BRC-1 chimera purification and E3 ubiquitin ligase assays**. (A) Purified chimeras visualized on stain-free gels (proteins do not run true to size as they were loaded on gel in sample buffer without heat denaturation). (B) Titration of E2 conjugating enzyme in E3 ubiquitin ligase assay shows a non-specific mono-Ub product in the absence of E2 enzyme (red). (C) Incorporation of mono- and di-HA-Ub into histone H2A as visualized by antibody against HA.

**S2 Fig. C-terminal GFP fusion to BRD-1 does not promote nuclear accumulation of the complex nor rescue of embryonic lethality in response to IR treatment**. (A) BRD-1 protein localization shown by direct GFP fluorescence in wild-type and mutant *brc-1* worms in respective germ line regions. PZ = proliferative zone; TZ = transition zone; EP = early pachytene; MP = mid pachytene; LP = late pachytene; DP = diplotene. Scale bar = 10μm. (B) Embryonic lethality of worms treated with 75Gys IR. C-terminal GFP fusion to BRD-1 did not rescue viability in the *brc-1* mutants. (C) Immunoblot (left) showing steady state levels of BRC-1 proteins from wild type and mutant whole worm extracts. Levels of mutant BRC-1 proteins normalized to wild type protein from three independent experiments (right).

**S3 Fig. Analysis of *brd-1(null)***. (A) BRD-1 exon structure and position of insertion of the stop-in cassette. Primer pairs (P1-P3) used for RT-PCR of wild type and *brd-1(null)* cDNA are indicated. P1 Forward: cgccacatttcaacagaaacc, P1 Reverse: gcttctttgctgtagtcgtg; P2 Forward: cgcgtaattcgacaaaacgc, P2 Reverse: gcattaataactgcacccgc; P3 Forward: ggctcaacattagaaacaacgc, P3 Reverse: gatcaataatgcacgctctcag. *ama-1* was used as control (86). B) No GFP fluorescence was observed in *brd-1(null)::gfp* worms. Scale bar = 10μm. (C) Male self-progeny (left Y axis; n=12) and embryonic lethality (right Y axis; n=18) of *brc-1(null)* and *brd-1(null)* worms.

**S4 Fig. Embryonic lethality of different fluorescent protein fusions and GFP::BRC-1 RING purification**. (A) Embryonic lethality in the presence of 75Gys IR was examined in *brd-1(null)* (n=21), *brc-1(null)* (n=21), *gfp::brd-1* (n=12), *brc-1(null) gfp::brd-1* (n=10), *brd-1::gfp* (n=11), *brc-1(null) brd-1::gfp* (n=18), *mScarlet::brc-1* (n=14), *gfp(nd)::brc-1* (n=14). (B) Direct GFP fluorescence of germ cell nuclei from *brc-1(null) gfp::brc-1*. Scale bar = 10μm. (C) Purified GFP::BRC-1 RING protein visualized on stain-free gel (protein does not run true to size as it was loaded on gel in sample buffer without heat denaturation). (D) Graph of relative steady state levels of GFP::BRC-1, mScarlet::BRC-1, and GFP^nd^::BRC-1 in the *brd-1(null)* mutant.

